# Alternative splicing of transposable elements in human breast cancer

**DOI:** 10.1101/2024.09.26.615242

**Authors:** Alex Nesta, Diogo F. T. Veiga, Jacques Banchereau, Olga Anczukow, Christine R. Beck

**Affiliations:** The Jackson Laboratory for Genomic Medicine, Farmington, CT 06032 USA; Department of Genetics and Genome Sciences, University of Connecticut Health Center, Farmington, CT 06030, USA; Department of Translational Medicine, School of Medical Sciences, University of Campinas (UNICAMP), Campinas, SP 13083, Brazil; Institute for Systems Genomics, University of Connecticut, Storrs, CT 06269, USA; Immunoledge LLC, Montclair, NJ, 07042, USA

## Abstract

Transposable elements (TEs) drive genome evolution and can affect gene expression through diverse mechanisms. In breast cancer, disrupted regulation of TE sequences may facilitate tumor-specific transcriptomic alterations. We examine 142,514 full-length isoforms derived from long-read RNA sequencing (LR-seq) of 30 breast samples to investigate the effects of TEs on the breast cancer transcriptome. Approximately half of these isoforms contain TE sequences, and these contribute to half of the novel annotated splice junctions. We quantify splicing of these LR-seq derived isoforms in 1,135 breast tumors from The Cancer Genome Atlas (TCGA) and 1,329 healthy tissue samples from the Genotype-Tissue Expression (GTEx), and find 300 TE-overlapping tumor-specific splicing events. Some splicing events are enriched in specific breast cancer subtypes – for example, a TE-driven transcription start site upstream of *ERBB2* in HER2+ tumors, and several TE-mediated splicing events are associated with patient survival and poor prognosis. The full-length sequences we capture with LR-seq reveal thousands of isoforms with signatures of RNA editing, including a novel isoform belonging to *RHOA*; a gene previously implicated in tumor progression. We utilize our full-length isoforms to discover polymorphic TE insertions that alter splicing and validate one of these events in breast cancer cell lines. Together, our results demonstrate the widespread effects of dysregulated TEs on breast cancer transcriptomes and highlight the advantages of long-read isoform sequencing for understanding TE biology. TE-derived isoforms may alter the expression of genes important in cancer and can potentially be used as novel, disease-specific therapeutic targets or biomarkers.

**One Sentence Summary:** Transposable elements generate alternative isoforms and alter post-transcriptional regulation in human breast cancer.

## Introduction

Transposable elements (TEs) comprise approximately half of the human genome (International Genome Sequencing Consortium *et al*., 2001) and play an important role in genomic regulation. Although most TEs in the human genome are no longer capable of retrotransposition due to accumulated mutations or host repressive mechanisms, many retain functional motifs that can impact gene expression and splicing in both normal and disease contexts (Percharde *et al*., 2018; McKerrow *et al*., 2022).

In some cancers, including breast tumors, TEs can alter gene expression through aberrant splicing (Zarnack *et al*., 2013; Jang *et al*., 2019; Clayton *et al*., 2020). Genome-wide methylation studies have shown that tumor-associated DNA demethylation occurs more frequently near TEs (Kong *et al*., 2019), and loss of DNA methylation can regulate TE-derived transcription start sites for oncogenes like *LIN28B* and *MET* (Miglio *et al*., 2018; Jang *et al*., 2019). Some of these events result in tumor-specific TE-chimeric antigens (Shah *et al*., 2023) and are emerging as an important source of neoantigens (Merlotti *et al*., 2023). The discovery and characterization of tumor-specific TE splicing events has been limited by short-read RNA sequencing (RNA-seq) (Sharon *et al*., 2013; Vaquero-Garcia *et al*., 2016).

We demonstrate the utility of long-read RNA sequencing (LR-seq) in studying the transcriptomic effects of TEs in cancer. Our work reveals the substantial contribution of TEs to novel isoforms and splice junctions in breast cancer. Some of these isoforms are highly prevalent across hundreds of patients in (TCGA) and show enrichment in specific breast cancer subtypes (Perou *et al*., 2000).

Our analysis of alternatively spliced TEs in breast cancer transcriptomes encompasses alternative first exons, cassette exons, and last exons. Our examination of the transcriptomic alterations of TEs extends to post-transcriptional RNA editing. This process is upregulated in breast cancer (Sagredo *et al*., 2020) and we detail isoform-specific RNA editing events in our data. Finally, we examine how polymorphic TEs can lead to unannotated splicing events, demonstrating the potential for long-read isoform analysis in personalized medicine.

This study enhances our understanding of TEs in the context of breast cancer and demonstrates the myriad ways that TEs can influence cancer transcriptomes. Our insights serve as a foundation for the development of new strategies to discover cancer-specific neoantigens for use in immunotherapies.

## Results

### LR-seq identifies novel TE-containing isoforms in human breast tumors

Using Pacific biosciences long-read RNA-sequencing, we previously mapped 142,514 full-length isoforms across 30 breast samples, including primary tumors and healthy tissues, patient-derived xenografts, and cell lines (Veiga *et al*., 2022). Here, to investigate the prevalence of TEs in human breast cancer, we intersect the exons from the breast-specific LR-seq isoforms from our previous study (Veiga *et al*., 2022) with LINE, SINE, DNA, and LTR annotations from RepeatMasker (hg38) (Smit, AFA, Hubley, R, and Green, P, 2013) (**Fig. 1A**). We find that approximately half (72,388 / 142,514) of our LR-seq-derived isoforms contain at least one exon that overlaps a TE, and only 24% of these TE-containing exons map to a known isoform from the GENCODE v30 reference transcriptome (**Fig 1B**).

**Figure 1.**
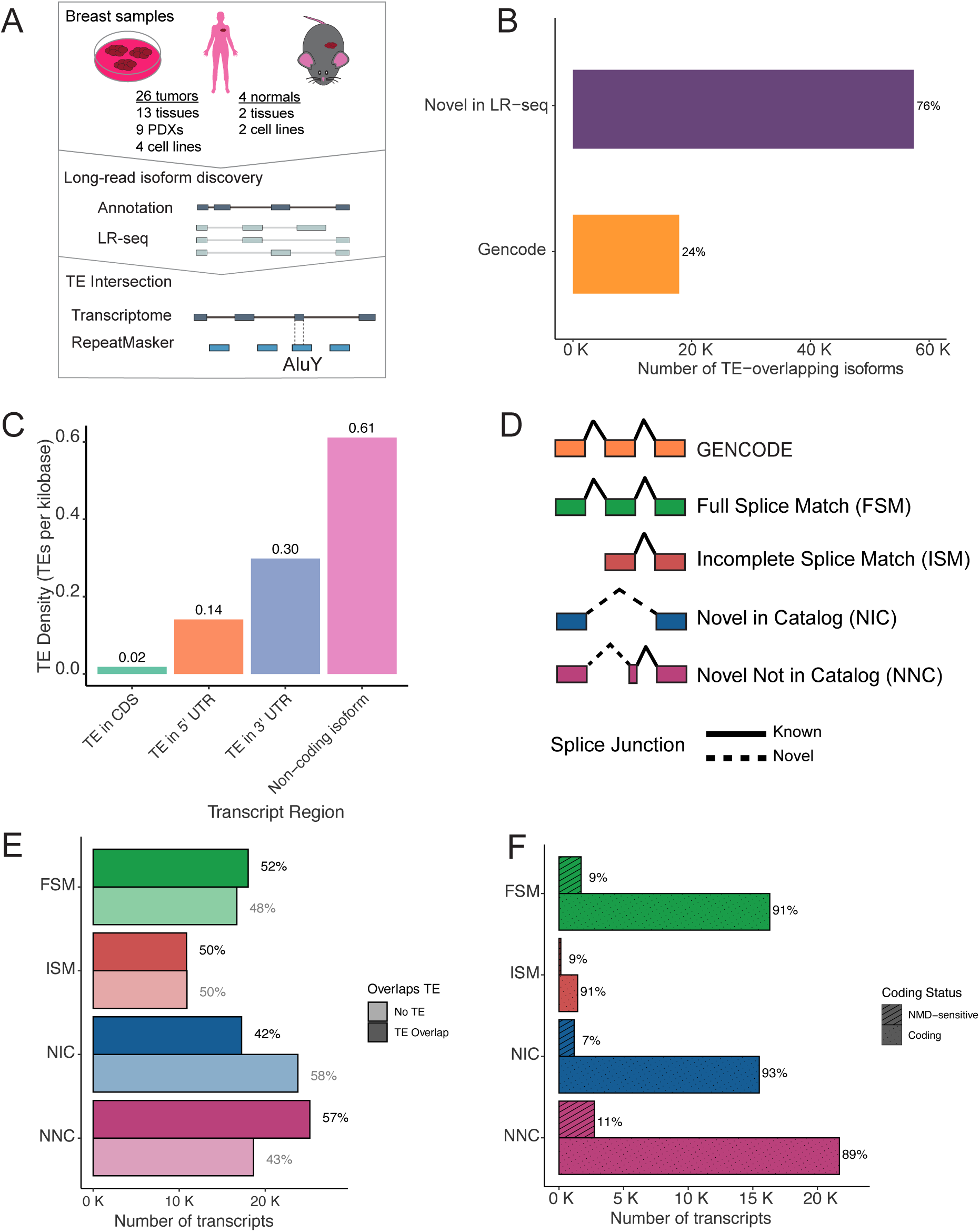
LR-seq identifies novel TE-containing isoforms in human breast tumors. **(A)** Overview of the study design and workflow. (1) Breast samples, including tumors, tissues, cell lines, and xenografts, are subjected to both long-read sequencing (LR-seq) and short-read RNA sequencing (RNA-seq). (2) Novel isoforms from LR-seq are compared to the GENCODE reference transcriptome. (3) TE-containing isoforms are found by intersecting LR-seq isoforms with transposable elements (TEs) from RepeatMasker. (**B**) Percentage of TE-containing isoforms that are in GENCODE (known isoforms) or are novel to the LR-seq dataset. (**C**) Normalized density of TE-overlapping exons across different regions of LR-seq isoforms: 3’ untranslated region (UTR), 5’ UTR, coding sequence (CDS), and non-coding regions. Non-coding isoforms lack an open reading frame prediction and may arise from aberrant splicing, pseudogenes or non-coding RNAs. (**D**) Schematic of SQANTI categories for classification of LR-seq isoforms based on splice junction alignment relative to GENCODE (Tardaguila *et al*., 2018). (**E**) Proportion of TE-containing isoforms within each SQANTI category. (**F**) Protein-coding potential and nonsense-mediated decay (NMD) sensitivity of LR-seq isoforms across SQANTI categories. NMD-sensitive isoforms contain premature termination codons >50 bp upstream of the final splice junction.

We next characterize the distribution of TEs in our LR-seq transcriptome by calculating their density per kilobase across different transcript regions (**Fig. 1C**). This analysis reveals a clear trend in TE density: coding sequences (CDS) show the lowest density (0.02 TEs/kb), followed by 5’ UTRs (0.14 TEs/kb), 3’ UTRs (0.30 TEs/kb), and non-coding isoforms (0.61 TEs/kb). The low TE density in CDS regions likely reflects strong selective pressure against disrupting protein-coding sequences. The next most TE-dense region was the 5’ UTR, where TEs can modulate translation by forming upstream open reading frames or be utilized in the initiation of transcription in tumors (Kitano, Kurasawa and Aizawa, 2018; Attig *et al*., 2019). The higher density in the 3’ UTRs supports previous findings that TEs in this region can affect mRNA stability, localization, and translational efficiency; moreover, the enrichment of TEs in the 3’UTR has been implicated in tumorigenesis (Fitzpatrick and Huang, 2012; Daniel, Lagergren and Öhman, 2015; Mayr, 2016; Chan *et al*., 2022; Gebrie, 2023).

To identify splicing categories of TE-containing isoforms, we next group them into categories based on their splice junction novelty *vs.* GENCODE using SQANTI (**Fig. 1D**) (Tardaguila *et al*., 2018). SQANTI classifies transcripts as Full Splice Match (FSM) if they match all splice junctions of a reference transcript, Incomplete Splice Match (ISM) if they match only some consecutive junctions, Novel in Catalog (NIC) if they contain new combinations of known splice sites, and Novel Not in Catalog (NNC) if they use at least one novel splice site. We observe 57% of the NNC isoforms overlapping with TEs (**Fig. 1E**). This represents the highest proportion of TE-containing isoforms among all SQANTI categories. Further analysis showed that ∼50% of the NNC novel splice junctions overlap a TE (18,316 / 43,400), highlighting the contribution of TEs to novel isoforms in cancer (**Fig. S1A**).

To determine if TE-overlapping LR-seq isoforms are likely to be degraded by nonsense-mediated decay (NMD), we next predict open reading frames from our isoforms. We find that the majority (>89%) of isoforms in each SQANTI category are potentially protein-coding and not NMD-sensitive (**Fig. 1F**). This observation is consistent with our previous findings (Veiga et al., 2022), where most isoforms, including those in the NNC category, contained a predicted open reading frame. To further support the coding potential of these TE-containing isoforms, we find perfect coding sequence matches for 5% of NNC isoforms with a TE in their predicted CDS using UniProt (**Fig. S1B**). Additionally, ribosome profiling data from nine breast cell lines (Vaklavas, Blume and Grizzle, 2020) supports the translation of ∼50% of NNC isoforms with a TE in their CDS (**Fig. S1C**). Taken together, our results provide compelling evidence that TEs contribute to a substantial portion of novel, potentially functional isoforms in breast cancer.

### LR-seq reveals preferential alternative splicing of TEs across the cancer genome atlas

We quantify the frequency and prevalence of TEs alternatively spliced in breast cancer with respect to normal breast tissue. We interrogate our previously published 310,000 alternative splicing (AS) events from LR-seq and GENCODE across TCGA (n = 1,135 breast tumors; 114 adjacent normal biopsies) and the GTEx (n = 1,329 samples across 12 tissues) datasets (Veiga *et al*., 2022) (**Fig. 2A**). To extract alternative splicing of TEs from our LR-seq transcriptome, we intersect the 5′ and 3′ ends of each AS event with LINEs, SINEs, DNA, and LTRs from RepeatMasker.

**Figure 2.**
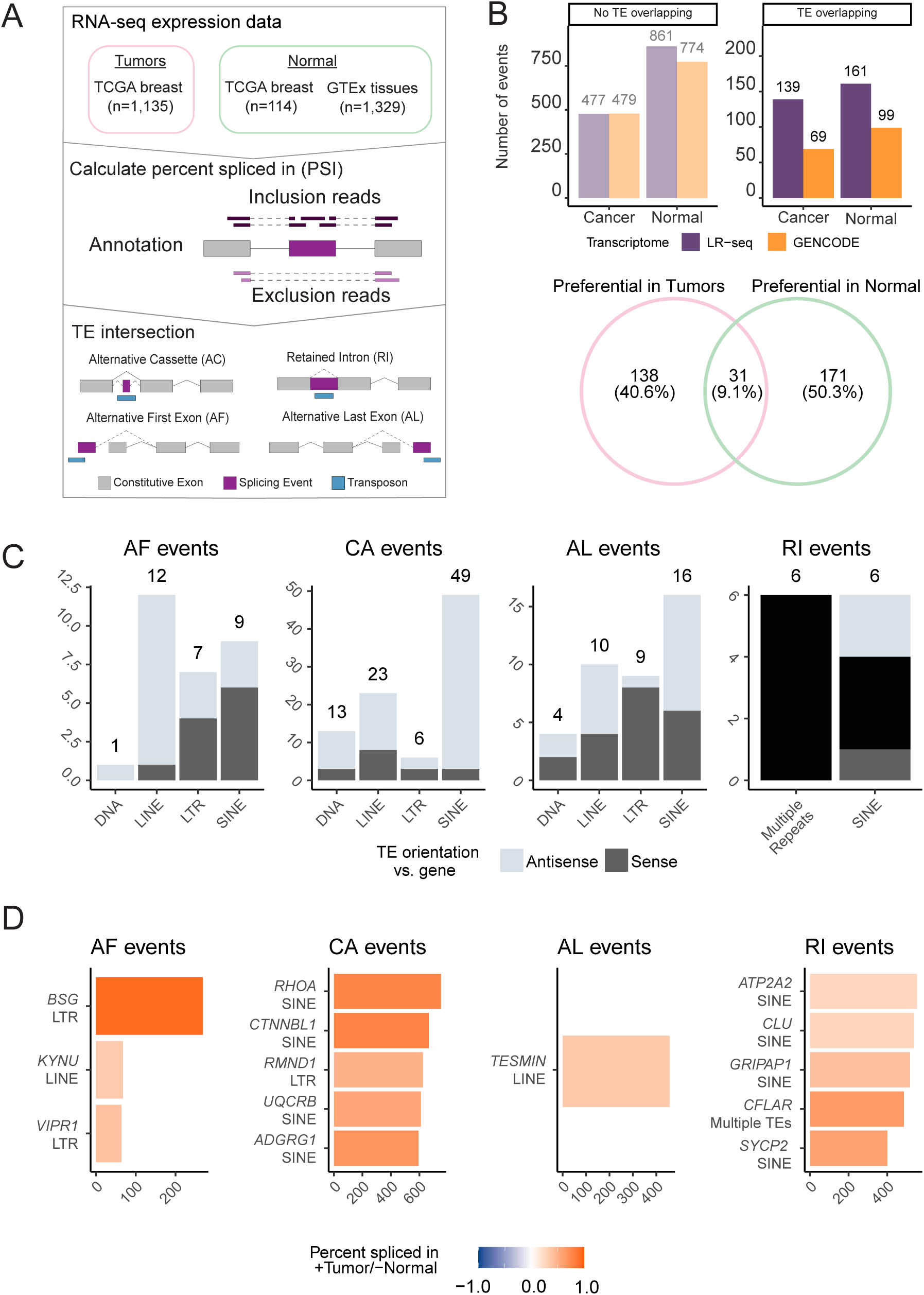
Some alternatively spliced TEs have higher expression in tumors. (**A**) Pipeline for identification of tumor specific AS events in TCGA breast tumors compared to normal breast tissues and GTEx tissues. (1) Breast cancer and normal tissue sample data were obtained from TCGA and GTEx for splicing quantification (Veiga *et al*., 2022). (2) Splicing was quantified as percent spliced in (PSI) using SUPPA2 (Trincado *et al*., 2018). PSI is calculated as (inclusion reads / [inclusion reads + skipping reads]). (3) Splicing events that overlapped TEs annotated in RepeatMasker are selected for further analysis. (**B**) Categorized AS events with a ΔPSI > 0.1 when comparing tumors to normal samples. Higher normal are events with ΔPSI < −0.1 and higher tumor events have a ΔPSI > 0.1. Events are further divided by transcriptome reference (LR-seq or GENCODE) and their overlap with a TE. The Venn diagram represents genes with TEs upregulated specifically in tumors or normal samples or shared between the two cohorts. (**C**) Distribution of TE classes and orientations for AS events with biased expression in tumors compared to normal tissues (see *methods*). Results are separated by alternative first (AF), cassette alternative (CA), alternative last (AL), and retained intron (RI) exons separated by TE class. TE orientation is represented with respect to gene orientation. (**D**) Quantification of canonical (GENCODE TSL1) to non-canonical (TE-containing LR-seq) isoform switching events for AF and CE events.

We identify 644 differential TE-overlapping AS events (20% of 3,095 total events) using SUPPA (Trincado *et al*., 2018), across TCGA Breast tumors and GTEx normal samples, defined by a difference in splicing by at least 10%. We find that 46% (300 / 644) of TE-mediated AS events are absent from Gencode. Furthermore, we observe that an approximately 44% - 56% (208 - 260) of TE-mediated AS events have biased usage in either breast tumors or normal tissues (**Fig. 2B**). The genes containing exonized TEs differ between tumor (171 genes) and normal (138 genes) tissues, with the two conditions sharing 31 genes (**Fig. 2B**).

To examine TE-mediated AS events with preferential splicing in tumors, we divide them by alternative first (AF), cassette alternative (CA), alternative last (AL), and Retained Intron (RI) events **(Fig. 2C)**. After categorizing these events by TE class and orientation, we find overrepresentation of TE classes with specific splicing mechanisms that have been detailed previously. For example, 12/13 LINE-overlapping alternative first exons initiate specifically in antisense-orientation LINEs. Previous reports showed that LINE-1 (L1) contains an antisense RNA polymerase II promoter capable of driving the expression of pro-tumorigenic coding and non-coding genes (Cruickshanks and Tufarelli, 2009; Criscione *et al*., 2016, 2016; Honda *et al*., 2020; Xu *et al*., 2023). We find that (3 / 13) antisense AS events align with an L1 antisense promoter consensus sequence (**Fig. S2A**) described in (Suzuki *et al*., 2002; Mätlik, Redik and Speek, 2006). Only (3 / 13) of our antisense L1 alternative first exon events were derived from HS, PA2, or PA3 elements (**Fig. S2B**), which are evolutionarily younger and more likely to contain an intact antisense promoter (Khan, Smit and Boissinot, 2006; Beck *et al*., 2010; Macia *et al*., 2011). We also observe a striking proportion of SINEs in our AS cassette exons that are in the antisense orientation (46 / 49); This observation is backed by a mechanism first observed by Sorek et al. 2002 (Sorek, Ast and Graur, 2002; Zarnack *et al*., 2013). We find AS antisense SINEs are largely Alu elements (40 / 46). These elements are overrepresented compared to all other intronic TEs in our dataset (chi-squared empirical *p*<9^-05^) (**Fig. S2C**). Furthermore, antisense SINEs serve as splice acceptors in our alternative last exons far more often than sense SINEs (23 / 25). Conversely, sense orientation SINEs are solely seen as alternative transcription termination sites in our LR-seq transcriptome (**Fig. S2D**).

Next, we investigate canonical to non-canonical isoform switching events that are enriched in breast cancer. To do this, we select the most frequent AF, CA and AL events in tumors and compare their TE-mediated splicing events with high-confidence (transcript support level 1) GENCODE annotated exons (**Fig. 2D**). We find three non-canonical TE-mediated AF events where LTRs or LINEs may increase transcription of the breast cancer associated genes *BSG*, *KYNU*, and *VIPR1* (Moody and Jensen, 2006; Ma *et al*., 2014; Liu *et al*., 2019). *BSG* is involved in tumor invasion and metastasis in multiple tumor types and methylation of its promoter is a proposed cancer prognostic biomarker (Fu *et al*., 2023), *KYNU* plays a role in immune regulation and is a proposed target for breast cancer metastasis (Girithar *et al*., 2023), and *VIPR1* is involved in cell proliferation and survival through its role in arginine metabolism (Fu *et al*., 2022). Two of our AF events are enriched for particular breast cancer subtypes: *BSG* in HER2+ and Luminal B breast cancers; *KYNU* in HER2+ (**Fig. 2D, S3A**). Isoform switches involving CE events are prevalent in hundreds of patients (>500 for the top 5 events). The most prevalent alternative CE we quantify (751 / 1,135 breast tumors) resides in the coding region of *RHOA,* a gene involved in migration, metastasis, and therapeutic resistance in breast cancer (Humphries, Wang and Yang, 2020). Additional isoform switches involving cassette exons include the genes *CTNNBL1, RMND1, UQCRB,* and *ADGRG1,* all of which have potential pro-tumorigenic roles or serve as biomarkers in breast cancer prognosis (Dunning *et al*., 2016; Kim *et al*., 2017; Li *et al*., 2017; Sasaki *et al*., 2021). Notably, the splicing events in *ADGRG1* and *CTNNBL1* are enriched in Luminal B samples (**Fig. 2D, S3A**). There was only one AL exon switch from canonical to non-canonical for the gene *TESMIN*. This gene is implicated in non-small cell lung cancer (Grzegrzolka *et al*., 2019) and contains an AL exon shared across >500 breast tumors without a clear subtype enrichment (**Fig. 2D, S3A**). In conclusion, we find that 208 TE-mediated AS events (171 unique genes) happen more frequently in breast tumors than in normal tissues and this difference in splicing is consistent across hundreds of tumor samples. Furthermore, we observe TE-mediated AS events in oncogenes that may play a role in tumorigenesis. However, the relationship between these AS events and gene expression is complex and requires further investigation. These findings underscore the importance of further investigating functional consequences of TE-mediated AS events in breast cancer and their potential as novel biomarkers or therapeutic targets.

### Alternative splicing of TEs is breast cancer subtype specific and associates with patient survival

We previously identified AS events enriched in one of four breast cancer subtypes: Luminal A, Luminal B, HER2 positive, and basal, and hypothesized that TEs may be alternatively spliced in a subtype-specific context (Veiga *et al*., 2022). By intersecting subtype-specific AS events with our catalog of TE-mediated AS events, we found 67 subtype-enriched events (p < 0.05) across 55 genes **(Fig. S3A)**. The most enriched TE splicing events included an LTR alternative first exon in the *AP2A2* gene found in 61 basal breast tumors, and an antisense Alu element in the *ERBB2* oncogene in the in 12 HER2+ tumors. *AP2A2* encodes a transcription factor that regulates the tumor suppressor DLEC1 and is a putative breast cancer target (Niranjan *et al*., 2023). *ERBB2*, which is frequently amplified and overexpressed in HER2+ breast cancer (Liu *et al*., 1992), harbored multiple *Alu* exonization events (Jang *et al*., 2019). In the luminal A subtype, we observed enrichment of an Alu exon in the long non-coding RNA *CASC2*, which is a reported tumor suppressor (Zhang *et al*., 2019). The luminal B subtype exhibited exonization of LINE1 elements in the mitotic kinase *AURKA*, an oncogene involved in enhancing stem-like features in breast tumors (Zheng *et al*., 2016).

To investigate the potential clinical relevance of these TE-overlapping AS events, we examined whether they were associated with patient survival (Veiga *et al*., 2022). The AF exon overlapping an LTR transposon in the *AP2A2* gene described above was enriched in 61 basal-like tumors (**Fig. S3B**) and associated with poor prognosis (p < 0.0013). Additionally, two TE-mediated AS events that exonized LINE1 elements in the *DUXAP9* and *ECHDC1* genes were also linked to unfavorable survival outcomes (**Fig. S3B**). Overexpression of the *DUXAP9* pseudogene is a prognostic biomarker in renal cell carcinoma (Chen *et al*., 2019) and the *ECHDC1* tumor suppressor may be disrupted by the LINE insertions (Jaiswal *et al*., 2021). These findings suggest subtype-specific TE splicing events may influence patient outcomes, particularly for more aggressive basal-like and HER2+ breast cancer subtypes.

### LR-seq captures ADAR editing in full-length transcripts

A 3′ UTR can span the last exon of a gene, and can contain multiple complementary pairs of *Alu-*SINE TEs that serve as substrates for RNA editing by Adenosine Deaminases acting on RNA or ADAR enzymes (Kim *et al*., 2004; Levanon *et al*., 2004; Sagredo *et al*., 2018). ADAR expression and editing are upregulated in breast cancer (Sagredo *et al*., 2020). The repetitive nature of *Alu* elements makes them challenging to study in the context of ADAR editing due to high sequence similarity compounded by editing-induced mismatches (Liu *et al*., 2014). LR-seq presents a unique opportunity to examine ADAR editing in the context, highly-accurate, full-length isoforms (Sharon *et al*., 2013; Liu *et al*., 2023). To identify ADAR edits captured with LR-seq, we used REDItools (Picardi and Pesole, 2013) to identify A > G mismatches which are the sequenced product of A > I deamination events (**Fig. 3A**). Since ADAR editing is most prominent in the 3′ UTR of transcripts (Levanon *et al*., 2004), we intersected putative edits with TE-containing last exons annotated in our LR-seq transcriptome. A > G and complementary T > C substitutions occurred most frequently in TEs versus non-TE regions (odds ratio > 1.0, **Fig. 3B**), and these edits primarily resided in *Alu-*SINE TEs (**Fig. 3C**). Identifying ADAR editing signatures in the last exons of our LR-seq isoforms revealed thousands of isoforms in both known and novel SQANTI categories (**Fig. 3D**). The identification of these events in so many novel isoforms highlights the prevalence of ADAR editing in breast cancer and suggests these events may have been missed in short read studies. To prioritize tumor-relevant ADAR events, we overlapped them with AL exon events enriched in breast tumors compared to normal tissues (**Fig. 3E**). The most prevalent event was found in over 1,000 breast tumors and resides in *TMED4*, where ADAR editing is a proposed prognostic marker for bladder cancer (Tang *et al*., 2023). LR-seq revealed T > C mismatches in complementary Alus within an extended 3’ UTR of TMED4 (**Fig. 3F**), indicating ADAR editing in breast cancer.

**Figure 3.**
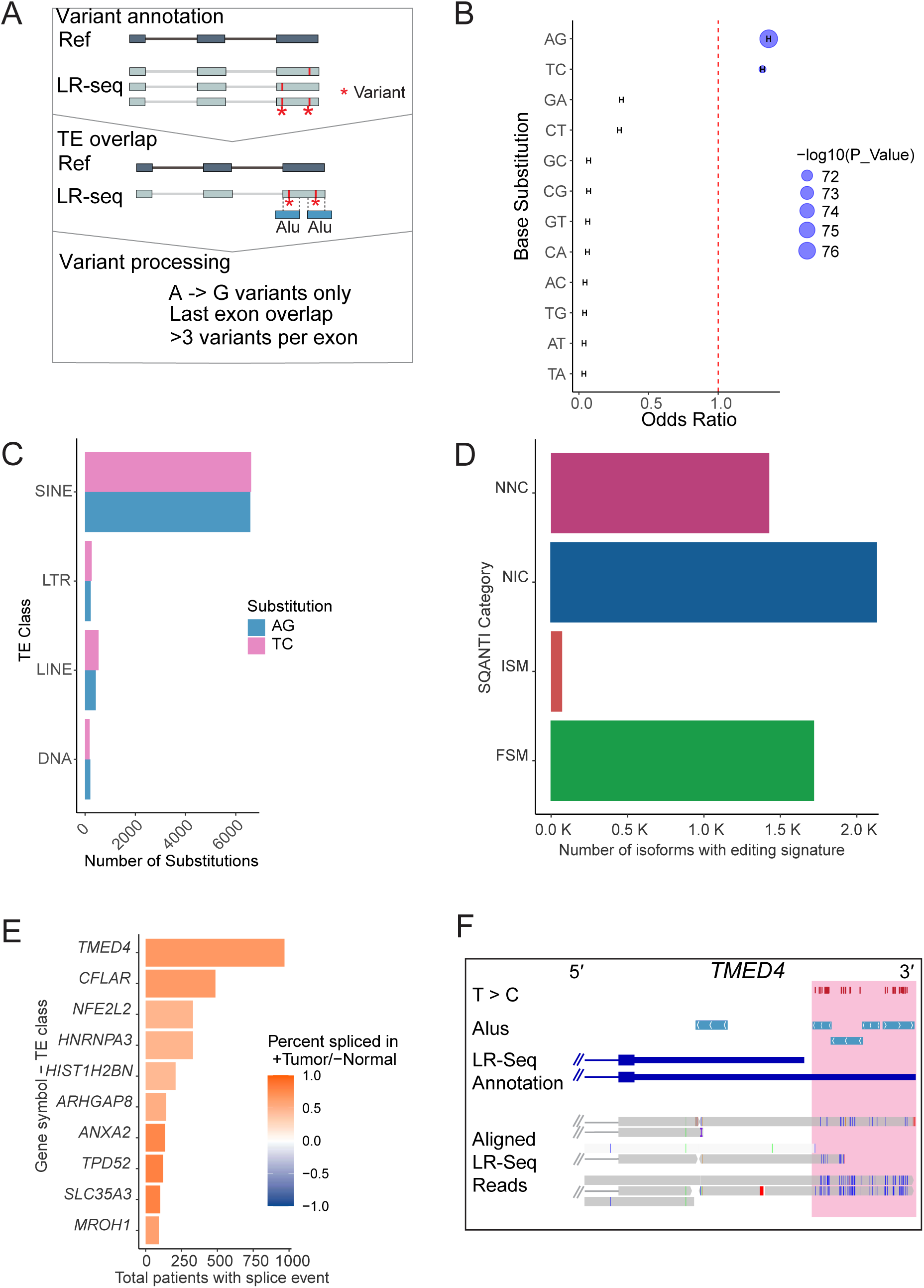
LR-seq detects breast cancer ADAR editing of *Alu* elements. (**A**) Pipeline to discover ADAR edits in LR-seq reads. (1) Mismatches against hg38 are determined using LR-seq reads with REDItools. (2) To identify mismatches/edits in TEs, these are intersected with TE annotations. (3) ADAR editing events are identified as A>G mismatches that meet specific criteria (see *methods*). (**B**) Calculated odds ratio and chi-squared test p-value of nucleotide substitutions in the last exons of LR-seq isoforms. The odds ratio measures the likelihood of a substitution occurring in a TE class versus outside a TE region; a ratio > 1 indicates higher likelihood. (**C**) Number of nucleotide substitutions in the last exons of LR-seq isoforms that overlap a TE class. (**D**) Isoforms containing at least 6 A>G mismatches in their last exon that overlap complementary *Alu* pairs. (**E**) Most common TCGA breast tumor AL events that overlap an ADAR editing signature. (**F**) Genome browser view of the TMED4 gene showing the T>C mismatch tracks between LR-seq reads and the reference genome, indicating ADAR editing sites. The Alu track shows sense and antisense Alu elements in the extended last exon. The annotated isoforms include the canonical TMED4 and a novel LR-seq isoform with the extended Alu-containing last exon. LR-seq read alignments highlight the T>C mismatches indicative of ADAR editing in this region.

As ADAR can edit within intronic sequences, we expanded our ADAR editing investigation beyond last exons (Tang *et al*., 2020). We find six LR-seq ADAR signatures overlapping the 300 tumor-enriched AS events we quantified in TCGA (**Fig. S4A**). This included an ADAR-edited Alu overlapping an CA exon in *RHOA*. LR-seq reads showed ADAR editing within and surrounding this alternatively spliced Alu element (Chen *et al*., 2023) (**Fig. S4B**), which is present in RHOA’s coding sequence. *RHOA* encodes a GTPase that is part of the RAS homolog family member A, and overexpression in breast cancer is a marker for tumor progression. (Bellizzi *et al*., 2008; Chan *et al*., 2010; Cheng *et al*., 2021). Comparison of the *RHOA* coding sequence with and without the Alu exon (UniProt C9JX21 vs P61586) revealed a misalignment after amino acid 138 (**Fig. S4C**). This *Alu-*containing isoform lacks a GTP binding domain found at positions 160-162 of the canonical sequence (**Fig. S4D**).

Taken together, these results demonstrate the ability of LR-seq to detect ADAR editing in full-length isoforms with potential cancer relevance, as exemplified in *TMED4* and *RHOA*.

### Polymorphic TE insertions can drive AS and are discoverable with LR-seq

Almost 10% of structural variants in the human genome result from TE insertions (Xing *et al*., 2009; Ebert *et al*., 2021). These polymorphic TEs are absent in the GRCh38 reference and typically overlooked in most RNA-seq studies. Previous studies combined both whole-genome sequencing and RNA-seq and discovered polymorphic TE insertions that impact gene expression (Cao *et al*., 2020). LR-seq can capture transcripts containing polymorphic TEs, but these read segments are generally discarded during genome alignment.

To investigate polymorphic TE insertions, we extracted LR-seq reads containing clipped, inserted, or deleted segments (≥25 bp) that failed to align to the reference (**Fig. 4A**); these sequences may map to structural variants between humans, including TEs. Across all 30 LR-seq samples, we identified ∼58,000 full-length, circular consensus reads that contain clipped and/or inserted unaligned segments (**Fig. 4B**). We performed homology searches against a TE sequence database to determine if the unaligned regions represented TE sequences (Storer *et al*., 2021) (**Fig. 4C**).

**Figure 4.**
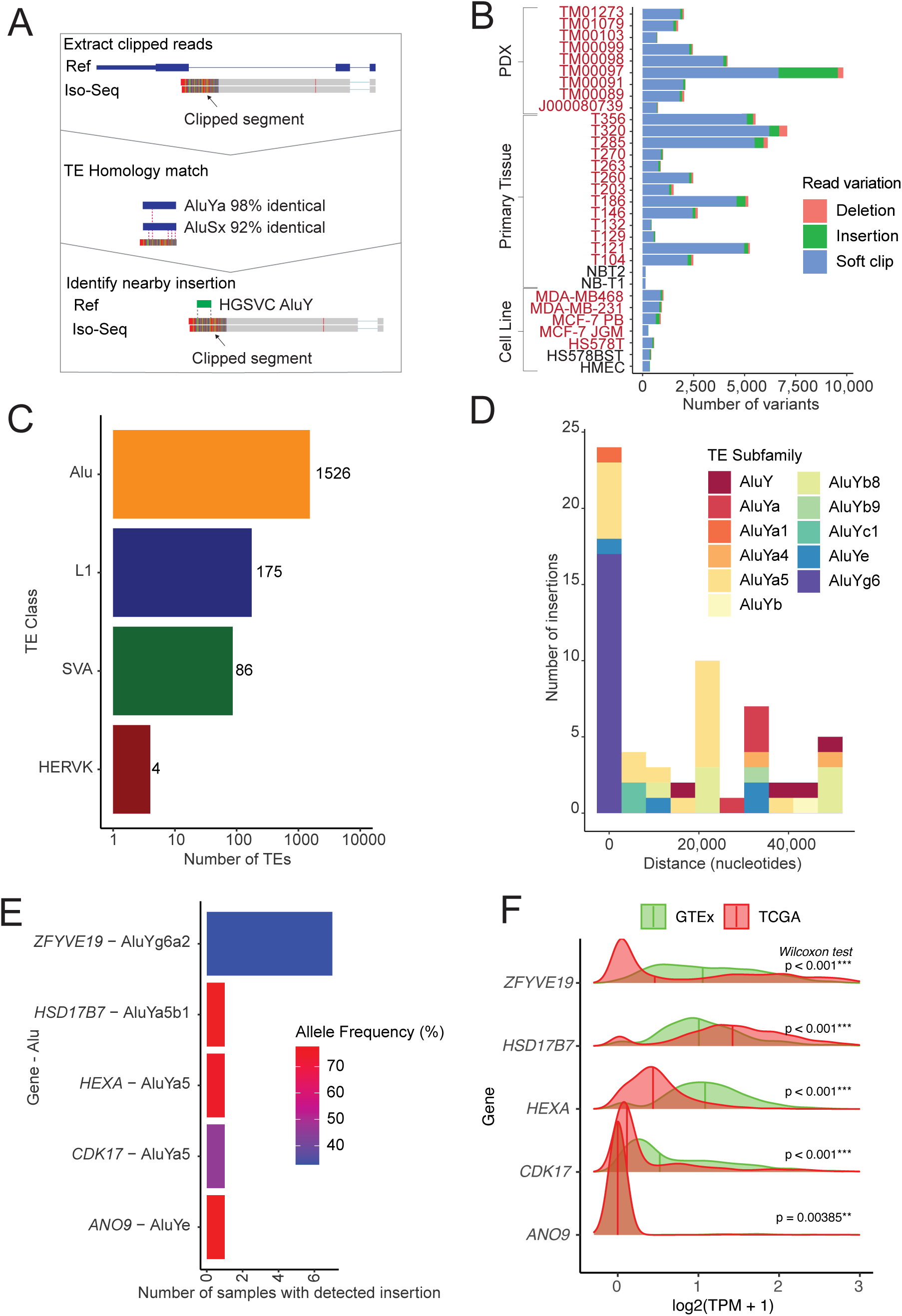
Identification of polymorphic TE insertions with LR-seq. **(A)** Pipeline to identify polymorphic TE insertions in LR-seq reads: (1) Extract LR-seq reads with clipped, deleted, or inserted segments that do not align to the reference genome. (2) Perform a homology search to identify TE sequences in the clipped region. (3) Intersect clipped TE segments with known polymorphic TE insertions from the HGSVC dataset (e.g., an Alu as depicted). (**B**) Number of LR-seq reads containing clipped/inserted/deleted segments across all LR-seq samples, separated into tumor (red) and normal (black) samples. The bars show reads with segments homologous to TE consensus sequence. (**C**) Breakdown of the highest homology scoring TE family matches in the clipped segments of LR-seq reads. (**D**) Distances between HGSVC validated *Alu* insertions and the LR-seq soft clipped regions of genes <50 kbp away. (**E**) Allele frequencies across 64 human genomes for the polymorphic TE insertion events identified from the soft-clipped LR-seq reads. (**F**) Expression levels across TCGA tumors and GTEx normal samples for genes containing an alternatively spliced polymorphic Alu detected by the LR-seq soft-clipped read analysis; the event in *ZFYVE19* is validated in **Fig. S6**.

Next, we intersected the genomic coordinates of the clipped segments with nearby (<50 kbp away) polymorphic TE insertions annotated in a set of 64 diverse human genomes from the Human Genome Structural Variation Consortium (HGSVC) (Ebert *et al*., 2021). We focused our attention on Alu subfamilies since they are the most active TE in the human genome. Our analysis revealed matching events between the LR-seq and HGSVC datasets (e.g., AluYa) and confirmed five alternative splicing events involving polymorphic Alu insertions (**Fig. 4D)** in the following genes *ANO9* (AL), *HSD17B7* (AF), *HEXA* (AL), *ZFYVE19* (AF), and *CDK17* (AL) (**Fig. 4E**).

We find these genes are differentially expressed in breast tumors *vs.* GTEx normal tissues. For example, *HSD17B7* and *ZFYVE19* contain AF polymorphic Alus, and showed increased tumor expression. *HEXA*, *CDK17*, and *ANO9* contain AL polymorphic Alus and had lower tumor expression overall (**Fig. 4F**).

We validated one of the AF events, a polymorphic AluY insertion in *ZFYVE19*, at both the genomic DNA level (confirming the insertion) and by amplifying the novel isoform containing the AluY-derived exon in MCF-7 cells versus a control cell line lacking the insertion (**Fig. S5A, B**). *ZFYVE19* has reported roles in cell cycle and immune regulation with relevance to cancer (Marina *et al*., 2010; Mandato *et al*., 2021; Bartolomé *et al*., 2023).

For *HSD17B7*, we identified an LR-seq read with a soft-clipped segment exhibiting high sequence similarity (96%) to a polymorphic *Alu*Y insertion from HGSVC, annotated as a putative alternative transcription start site (Fig. S5C). This *Alu*Y is a common human polymorphism (78% allele frequency) that may impact *HSD17B7* expression, a gene implicated in estrogen-driven breast cancer cell growth (Shehu *et al*., 2011).

In summary, our analyses uncovered several examples where polymorphic TE insertions generate alternative isoforms, highlighting the ability of LR-seq to comprehensively capture events missed by conventional short-read RNA-seq pipelines. Such findings have implications for understanding transcriptome and proteome diversity associated with disease.

## Discussion

LR-seq enabled our investigation into the transcriptional and post-transcriptional effects of TEs, and with these data, we find that TEs are a prevalent source of alternative splicing in breast cancer. TE-containing isoforms comprise more than half of the novel isoforms in our dataset and contribute to RNA-editing of thousands of isoforms that require further study. Some of these RNA-edited and TE-containing isoforms may be relevant for cancer progression or prognosis, and our data will serve as a thorough characterization of the consequences of TEs on breast cancer transcriptomes.

Our analysis reveals 300 preferentially spliced TEs common to hundreds of breast cancer patients that are rarely included in isoforms in normal tissues. Most TE-mediated splicing events in cancer occur in AF, CA, and AL events. AF events commonly result from alternatively spliced LINEs, which contain both forward and reverse RNA polymerase II promoters (Speek, 2001). CA splicing events largely consist of antisense SINEs, as *Alu*s contain a poly(A) tail which in antisense orientation acts as a poly(T) polypyrimidine tract that acts as a site for spliceosomal assembly and leads to the use of a downstream splice acceptor (Sorek, Ast and Graur, 2002). AL events contained LINEs and SINEs with poly(A) tails that include transcription termination sites (Chen, Ara and Gautheret, 2009).

Some TE-mediated splicing events in breast cancer were prevalent, enriched in specific subtypes, and associated with patient survival. We identified 67 subtype-enriched TE splicing events across 55 genes, including oncogenes and tumor suppressors. An LTR-driven alternative first exon in *AP2A2*, implicated in hematopoietic stem cell self-renewal (Ting *et al*., 2012), was enriched in basal tumors, while multiple *Alu* AF events were observed in HER2-positive subtype tumors in the *ERBB2* oncogene. Importantly, some of these subtype-specific TE splicing events, such as those in *AP2A2*, *DUXAP9*, and *ECHDC1*, were associated with patient survival, highlighting the potential of TE-mediated splicing as a source of novel biomarkers and therapeutic targets in breast cancer.

Tumor-specific splice junctions between coding exons and TEs can generate immunogenic peptides and elicit CD8+ T cell responses in patients with non-small cell lung cancer (Kong *et al*., 2019; Merlotti *et al*., 2023; Shah *et al*., 2023). The histone methyltransferase SETDB1 regulates the expression of several immunogenic exon-TE splicing junctions in a mouse model of lung cancer (Burbage *et al*., 2023). Our work has identified hundreds of TE-mediated splicing events enriched in breast tumors compared to normal tissues, laying the groundwork for future studies exploring the immunogenic potential, mechanisms of impact on patients, and potential therapeutic targets caused by aberrant splicing in breast cancer.

We also find novel ADAR editing sites in breast cancer that are not annotated in existing databases (Picardi *et al*., 2017). LR-seq enables isoform-specific ADAR edit identification, revealing AS events with preferential splicing in hundreds of tumors. One ADAR-edited isoform of *RHOA* may result from an ADAR-induced splice acceptor; *RHOA* has been implicated in lung cancer progression. RNA-editing is primarily thought to regulate splicing by modifying the binding sites of splicing factors (Lev-Maor *et al*., 2007; Solomon *et al*., 2013).

Polymorphic TEs in clipped isoform reads define a new way that TE effects on transcription can be identified. Previous studies required both whole genome sequencing and RNA-sequencing data to associate polymorphic TEs with gene expression and splicing alterations (Cao *et al*., 2020). Using LR-seq alone, we were able to reference existing databases of polymorphic TE insertions (Ebert *et al*., 2021) to confirm sequence homology with LR-seq reads. We anticipate that our method can be used to identify alternative splicing of somatic TE insertions in other contexts, expanding our understanding of TE-mediated transcriptomic diversity.

In conclusion, we demonstrate how TEs effect breast cancer transcriptomes, laying a foundation for future mechanistic studies on TE-mediated splicing in cancer. TEs influence transcriptional and post-transcriptional processes with significant disease implications. Our LR-seq dataset revealed disease-relevant alterations that were missed with traditional RNA-Seq and GENCODE references alone. Our findings provide insight into the impact of TE regulation on transcriptome fidelity in the context of breast cancer, leading to new avenues for diagnosis and treatment.

## Materials and Methods

### Generation of an LR-seq Transcriptome

We used LR-seq isoforms from the LR-seq QC-pass breast cancer transcriptome (Veiga *et al*., 2022). Briefly, 30 breast samples were sequenced with LR-seq and short-read RNA sequencing (RNA-seq). LR-seq data were processed using the ToFu pipeline obtained from (https://github.com/PacificBiosciences/IsoSeq_SA3nUP/wiki). Full length transcripts from 30 samples were merged with chain_samples.py from (https://github.com/Magdoll/cDNA_Cupcake) to create a baseline transcriptome annotation. The annotation was processed using SQANTI2 (Tardaguila *et al*., 2018), and QC-pass isoforms were selected if >10 SR-seq reads aligned to all splice junctions of LR-seq reads. For more information on LR-seq processing, please see (https://github.com/TheJacksonLaboratory/BRCA-LRseq-pipeline).

### Identification of TEs in the LR-seq transcriptome

Our QC-pass LR-seq transcriptome and the UCSC RepeatMasker annotation were loaded into an R session using the rtracklayer R package (Lawrence, Gentleman, and Carey, 2009). Exons were extracted from the LR-seq transcriptome. TE overlaps with LR-seq isoforms were identified using the find_overlaps function from the plyranges R package (Lee, Cook and Lawrence, 2019). See the supplemental script Figure_1.Rmd.

### Ribosome profiling support

We utilized isoform-level ribosome profiling from our previous study (Veiga *et al*., 2022) of LR-seq predicted open reading frames (ORFs) using ORQAS (Reixachs-Solé *et al*., 2020) and Ribosome profiling data for nine breast cancer cell lines data from (Vaklavas, Blume and Grizzle, 2020). ORFs were considered translated if their periodicity and uniformity scores reached the threshold of that for single-ORF housekeeping genes. We extracted transcript identifiers from isoform-specific ribosome profiling results and annotated TE-containing transcripts identified in our LR-seq transcriptome (Veiga *et al*., 2022). See the supplemental script Figure_1.Rmd.

### Uniprot Support

We previously predicted ORFs from LR-seq transcripts (Veiga *et al*., 2020) using TransDecoder (https://github.com/TransDecoder/TransDecoder) and aligned the ORFs to with UniProt annotations (UniProt Consortium, 2021). We compared transcript identifiers of LR-seq predicted ORFs with 100% UniProt identity match with TE-containing transcripts in our LR-seq transcriptome. See the supplemental script Figure_1.Rmd.

### Identification of AS TEs from SUPPA results

AS results from TCGA and GTEx using our LR-seq transcriptome were obtained (Veiga *et al*., 2022). The RepeatMasker annotation was obtained from the UCSC table browser and loaded into an R session using the tidyverse read_tsv function. Splice junctions were parsed into discrete chromosome, start, and end columns and converted into a genomic ranges object using the makeGrangesFromDataFrame function from the GenomicRanges R package. Five prime and three prime ends of each splicing event were extracted into separate ranges objects and intersected with RepeatMasker using the find_overlaps from the plyranges R package. See the supplemental script Figure_2.Rmd.

### Identification of ADAR-edits in exons

LR-seq reads were aligned to hg38 using minimap2 with options “-ax splice:hq-uf” and converted to bam format using samtools. We used REDITOOLS (Picardi and Pesole, 2013) REDItoolDenovo.py to identify substitutions against hg38 with option “-c 1”. REDITOOLS FilterTable.py was used to select substitutions that overlapped repeats and LR-seq transcriptome exons. Finally, overlapping repeat class and exon was annotated using RepeatMasker and REDITOOLS AnnotateTable.py. To count edits in AS exons, we loaded our SUPPA splicing quantification and REDITOOLS edits into an R session. AS exons were extracted from our TE_splice using plyranges, and overlaps were counted using the find_overlaps function. See script the supplemental script Figure_3.Rmd.

### Identification of polymorphic TEs

LR-seq reads were aligned to hg38 using minimap2 (Li, 2018) with options “-ax splice:hq-uf”. Using GMAP (Wu and Watanabe, 2005) incorrectly assigned clipped segments to distant >200 kbp away repeats despite specifying a cutoff. The python package pysam (https://github.com/pysam-developers/pysam) was used to extract clipped segments from LR-seq reads with >2 mapped exons. See script 4.1_extract_clipped_reads.py. nHMMER (Wheeler and Eddy, 2013) was used with Dfam 3.2 Transposable Element HMMs (https://www.dfam.org/releases/Dfam_3.3/families/Dfam_curatedonly.hmm.gz) (Storer *et al*., 2021). We extracted the top TE alignment for every clipped segment and assigned the read to a parent gene by intersecting neighboring exons from our LR-seq transcriptome using plyranges find_overlaps function. See the supplemental script Figure_4.Rmd. Insertions were intersected with HGSVC TE insertions using find_overlaps (Ebert *et al*., 2021).

## Declarations

### Ethics approval and consent to participate

This study utilized publicly available data from The Cancer Genome Atlas (TCGA) and the Genotype-Tissue Expression (GTEx) project. All samples in these databases were collected with patient consent and appropriate ethical approval from the relevant institutional review boards. Our study did not involve additional human participants, human data or tissue.

## Consent for publication

Not applicable. This manuscript does not contain data from any individual person.

## Acknowledgements

We thank members of the Beck lab including Parithi Balachandran, Ardian Ferraj, Peter Audano, and Eden Francoeur for critically reading this manuscript and specifically Parithi Balachandran for their support and insight in developing code and methodologies for analyses in the paper; Laura Urbanski of the Anczuków lab for cell line RNA used in PCR validations; Nathan Leclair, Mattia Brugiolo, and Brittany Angarola of the Anczuków lab for discussions and experimental protocols and insight. Elizabeth Tseng of Pacific Biosciences for her preliminary investigation of TEs in MCF-7 LR-seq data. The results in this paper are based on data generated by TCGA managed by the NCI and NHGRI. Additional information about TCGA is located at http://cancergenome.nih.gov. The data obtained from the GTEx Projects was supported by the Common Fund of the Office of the Director of the National Institutes of Health and more details are available at commonfund.nih.gov. Additional data in the manuscript were obtained from dbGap at www.ncbi.nlm.nih.gov/gap.

## Author Contributions

A.N. conceived and developed the methodology, performed bioinformatic analyses, wrote the manuscript, and performed experiments. D.F.T.V. performed bioinformatic analyses.

C.R.B acquired funding, advised in methodology development, provided expertise, and wrote the manuscript. O.A. and J.B. provided expertise and guidance for analyses.

## Availability of data and materials

The LR-seq and SR-seq data were acquired from our previous publication (Veiga *et al*., 2022) and are available at the European Genome Archive database (accession number EGAS00001004819). The source code, data inputs, and data outputs including supplementary tables are available from https://github.com/TheJacksonLaboratory/TE-LRseq-Analysis/ https://zenodo.org/records/13761416

---

**Figure S1.**
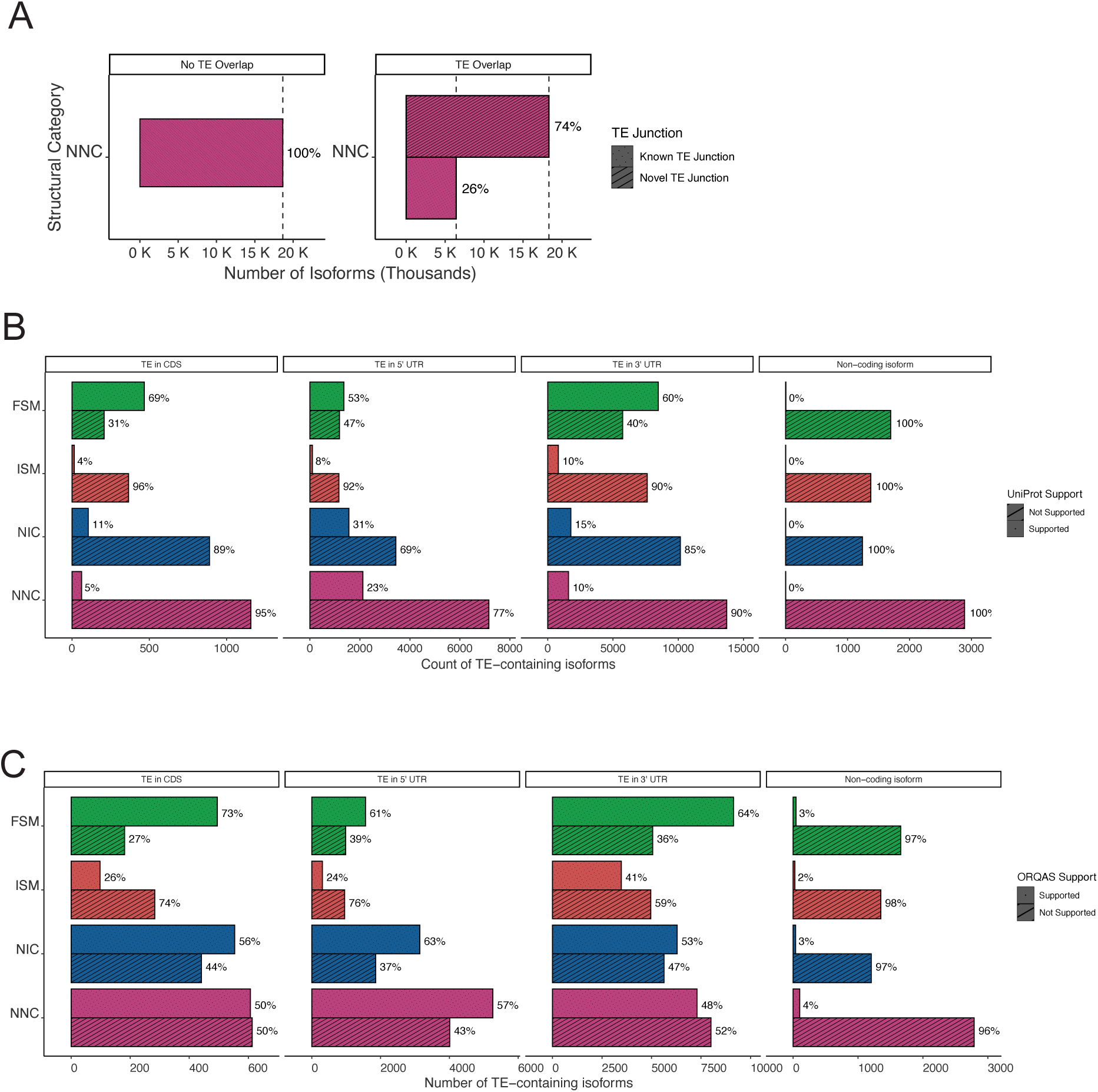
TE-containing isoforms are predicted to encode protein. (**A**) Proportion of NNC isoforms containing TEs. TE overlapping isoforms are further grouped by the presence of a novel splice junction present within a TE. (**B**) Protein-level support (100% coding sequence identity match) for TE-overlapping isoforms. Isoforms are divided by the location of the TE within a UTR or CDS. (**C**) Ribosome profiling support from 9 breast cancer cell lines as determined by ORQAS (Vaklavas, Blume and Grizzle, 2020). Isoforms are categorized as in panel **B**.

**Figure S2.**
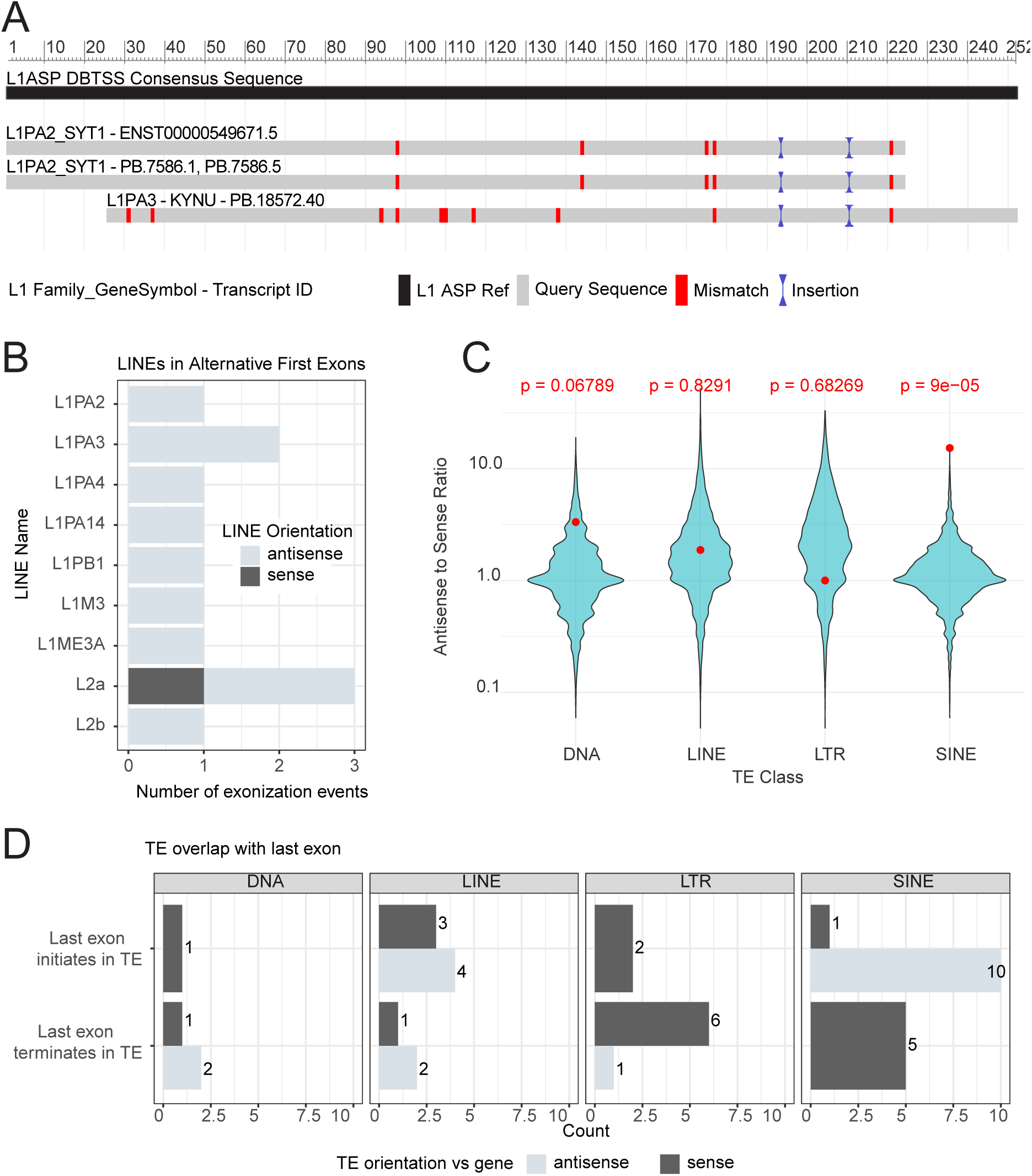
Quantitative examination of TE-mediated splicing mechanisms. (**A**) Alignment of the first 200bp of antisense L1-overlapping alternative first exons identified in the LR-seq dataset with an L1 antisense promoter consensus sequence. Matching segments are gray and the red indicates mismatches compared to the consensus L1 antisense promoter derived from the database of transcription start sties (DBTSS) (Yamashita *et al*., 2010). (**B**) Distribution of LINE subtypes among alternative first exons identified in the LR-seq dataset. LINE TEs are ordered on the Y axis in evolutionary order with L1PA2 being the youngest. TE orientation is annotated with respect to the parent gene’s orientation. (**C**) Ratios of antisense-to-sense oriented TEs within the intronic region of all GENCODE genes, separated by TE class. Red dots indicate the ratio of antisense-to-sense oriented cassette exons observed to be preferentially spliced in TCGA breast tumors compared to normal samples. (**D**) Count of alternative last exon events preferentially spliced in TCGA breast tumors categorized by whether the TE overlaps the splice acceptor or the transcription termination site of the last exon. Events are further separated by TE class and TE orientation with respect to the gene.

**Figure S3.**
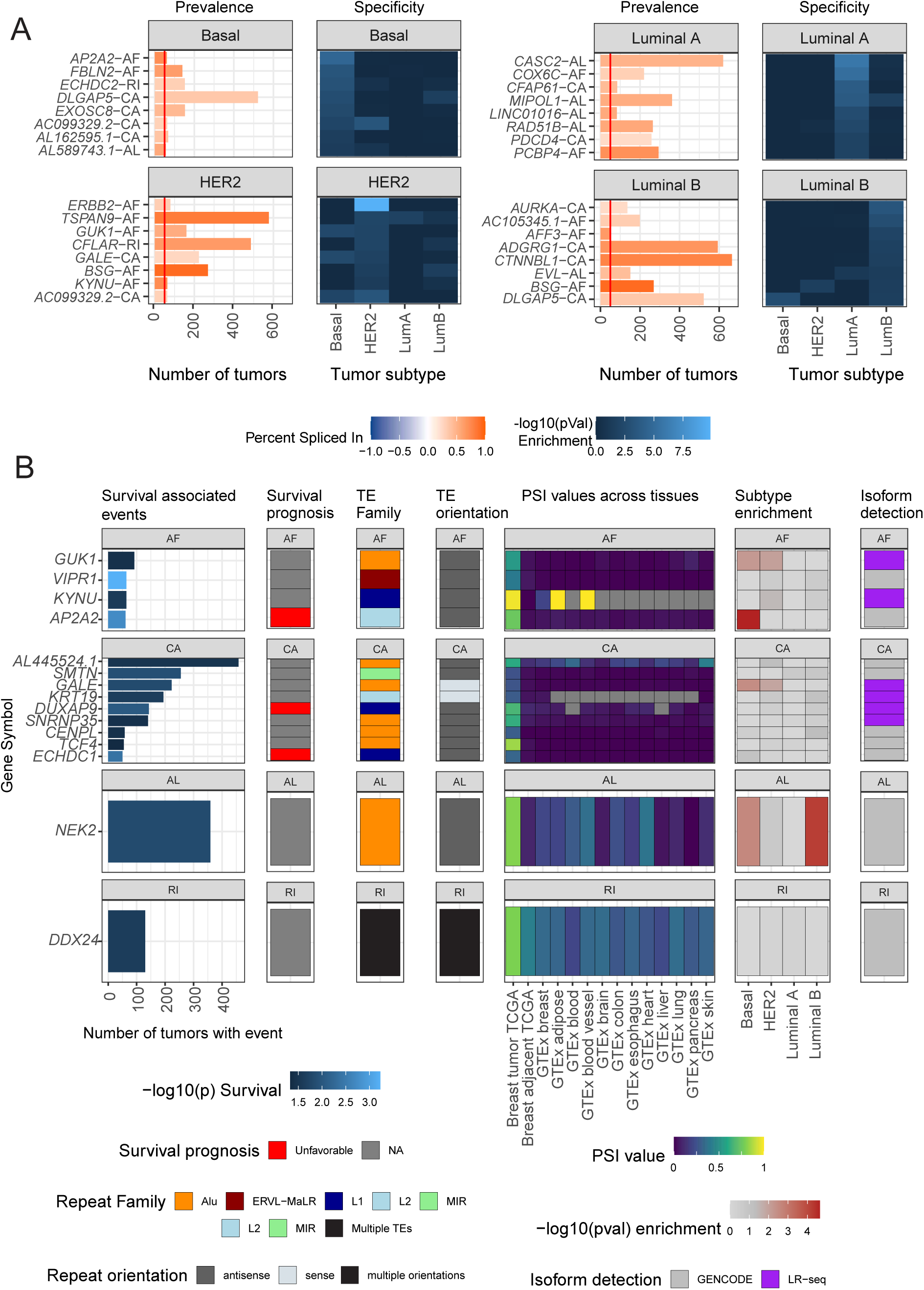
Alternatively spliced TEs are enriched in breast cancer subtypes and are associated with patient survival. **(A)** Top subtype-specific AS events (ordered by -log_10_(p) > 1.3) with TEs enriched in one of four breast cancer subtypes: Basal, HER2+, Luminal A, or Luminal B. Left panel: The number of tumors expressing each AS event, with the vertical red line indicating a minimum cutoff of 50 tumors. Events are labeled as “gene symbol - splicing event type” (AF: alternative first, CA: cassette alternative, AL: alternative last, RI: Retained Intron). Right panel: Heatmap showing the enrichment (-log_10_(p-value)) of each AS event across all subtypes. **(B)** TE-mediated alternative splicing events associated with patient survival, categorized by event type (AF, CA, AL, or RI). The heatmap columns represent the following information for each event: gene symbol, number of patients with the event, prognostic impact (favorable or unfavorable, if applicable), TE family, TE class, TE orientation relative to the gene, percent spliced in (PSI) values in TCGA and GTEx, breast cancer subtype enrichment, and the transcriptome (GENCODE or LR-seq) in which the AS event was annotated.

**Figure S4.**
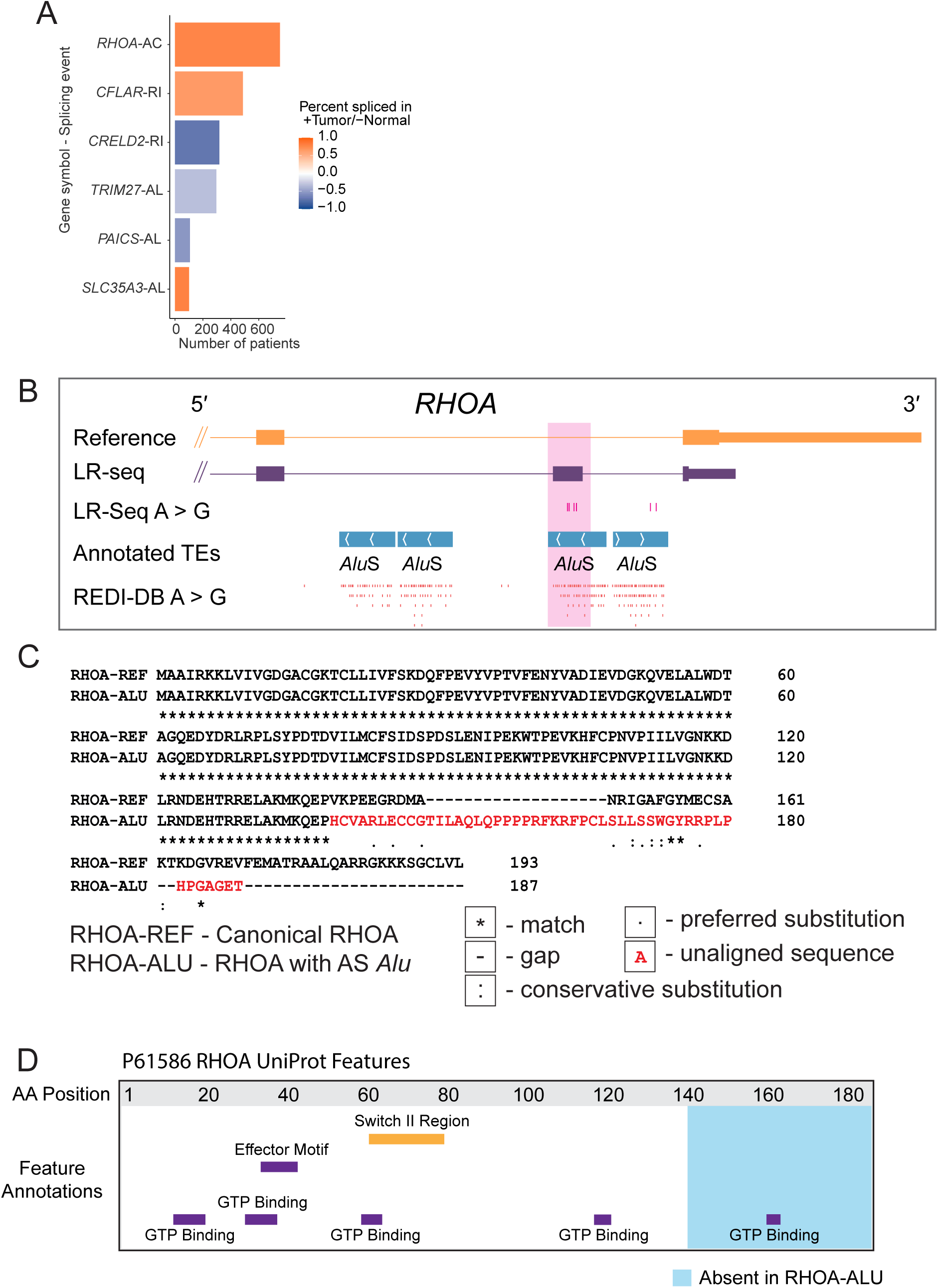
An ADAR-edited *Alu* disrupts the coding sequence of *RHOA.* **(A)** Top differential splicing events that also contain an ADAR editing signature detected from LR-seq reads; RI-Retained Intron, AC-Alternative Cassette, AL-Alternative Last. **(B)** Genome browser view showing that the RHOA gene contains an intronic Alu element with ADAR editing. Tracks shown include the GENCODE RHOA annotation, the LR-seq read with the Alu exonization event, mismatches highlighting the edited sites, annotated Alu pairs from RepeatMasker, and known ADAR editing sites from the REDI-DB database across GTEx samples. (**C**) Alignment comparing the reference RHOA protein coding sequence (UniProt P61586) to the predicted coding sequence when the Alu exon is included (UniProt C9JX21), showing misalignment after amino acid 138. (**D**) Protein domain architecture of the canonical RHOA isoform (P61586) from UniProt, highlighting the GTP binding domain from positions 160-162 that is disrupted in the Alu-containing C9JX21 isoform.

**Figure S5.**
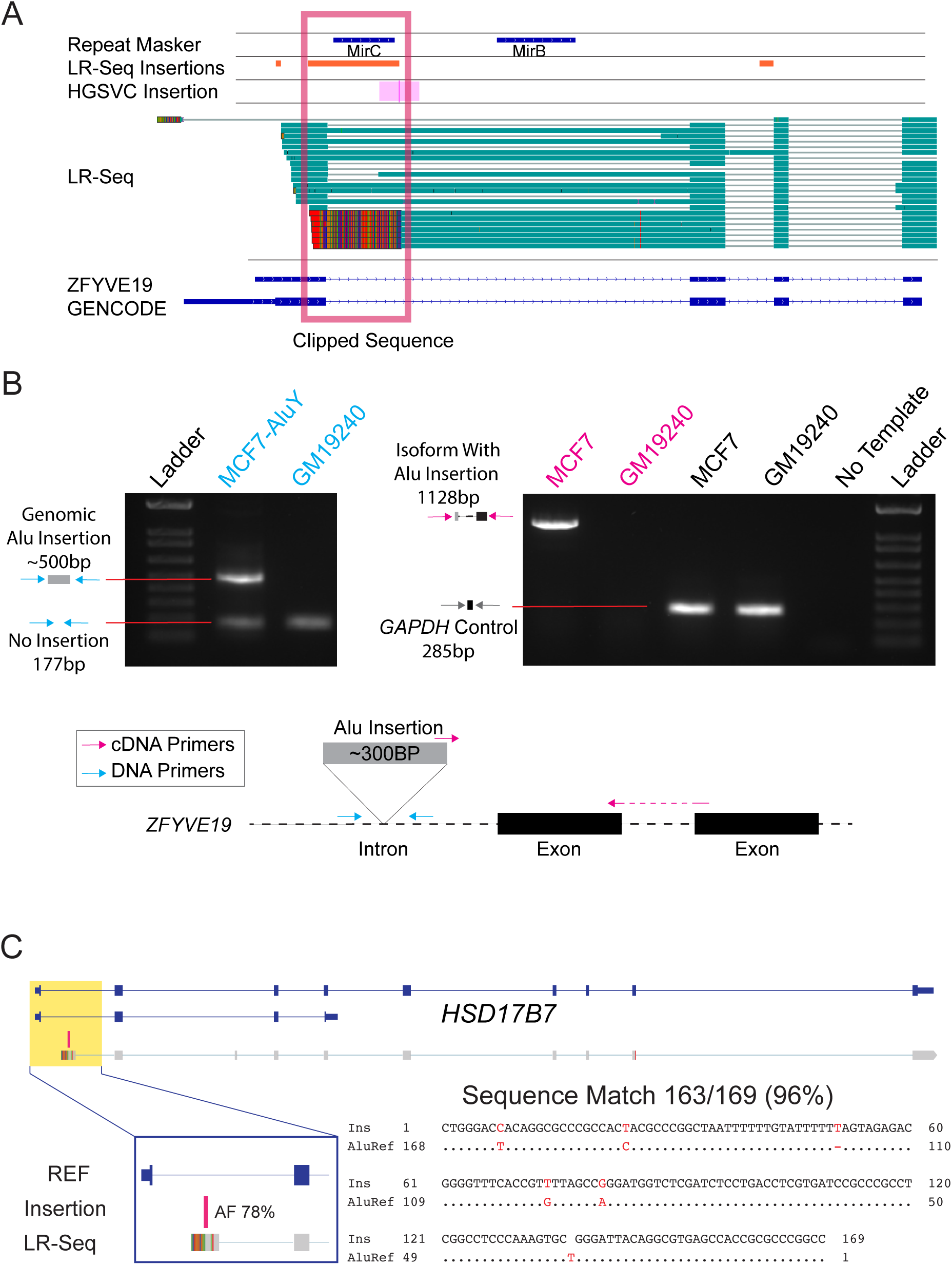
Validation of a polymorphic TE insertion in *ZFYVE19.* **(A)** Integrative Genomics Viewer image of a polymorphic *Alu*Y insertion (boxed region) annotated as an alternative first exon for *ZFYVE19*. (**B**) PCR validation of a polymorphic *Alu*Y insertion in *ZFYVE19* on the genomic level (DNA) and transcriptomic level (cDNA) in MCF-7 and GM19240 cell lines. Below the gel is a diagram of the two amplified regions of *ZFYVE19*. The blue arrows represent genomic DNA primers that flank the *Alu*Y insertion while the magenta arrows represent cDNA primers designed to span splice junctions that specifically amplify the isoform containing the *Alu*Y derived exon. (**C**) A polymorphic *Alu* is used as an alternative transcription start site for *HSD17B7*. The top of the figure contains two full-length isoforms for *HSD17B7* annotated in GENCODE and colored in blue. Below is an aligned LR-seq read containing a soft-clipping annotation at the 5′ end of the read. Above the soft-clipped annotation is a pink polymorphic TE insertion identified previously (Ebert et al., 2021) that has an allele frequency (AF) of 78%. Below is a magnified view of the soft-clipped segment of the read overlapping the polymorphic TE insertion. The soft-clipped segment of the read has 96% sequence homology with the *Alu* consensus sequence determined via Smith-Waterman alignment.

